# Reasons and Challenges of Adopting Flexitarian Diet: A Mixed-Methods Approach Using Natural Language Processing

**DOI:** 10.1101/2025.01.05.631383

**Authors:** Carla Djaine Teixeira, Djackson Garcia De Lima, Elias Jacob de Menezes Neto, Sávio Marcelino Gomes, Michelle Cristine Medeiros Jacob

**Affiliations:** Graduate Program in Social Sciences, Center for Human Sciences, Letters and Arts, Federal University of Rio Grande do Norte, Natal, RN, Brazil; Graduate Program in Nutrition, Federal University of Rio Grande do Norte, Natal, RN, Brazil; Institute of Digital Metropolis (IMD), Federal University of Rio Grande do Norte, Natal, RN, Brazil; Department of Nutrition, Federal University of Paraíba, João Pessoa, PB, Brazil; LabNutrir, Dpartment of Nutrition, University of Rio Grande do Norte, Natal, RN, Brazil

**Keywords:** flexitarianian diet, motivations, challenges, sociocultural factors, Brazil

## Abstract

In this study, we analyzed data from a large sample (n = 778) of individuals who identified as flexitarians, using Brazil as a case study. We applied Natural Language Processing (NLP) to characterize the reasons why individuals self-identify as flexitarians. We explored how these self-perceptions contribute to the construction of a flexitarian identity and identified the challenges this group faces in adopting a flexitarian diet. Our results indicate that there is no homogeneous flexitarian identity, but rather a diversity of categories, organized into five distinct clusters: ethical flexitarians (those who express greater concern for the environment and animals), flexible vegetarians (those who often express the desire to become vegetarian but still consume meat), meat lovers (those who enjoy meat but do not feel compelled to consume it), moderate consumers (those who refrain from meat on certain days of the week), and meat reducers (those who actively reduce their meat consumption). The latter three groups are more similar, as they tend to reduce their meat intake due to the belief that meat is not essential in their diet, although they still enjoy its taste. Additionally, we found that the main challenge faced by our participants is directly related to the social environment, specifically the difficulty in finding alternatives to meat, which makes the love for meat’s taste and meat-centric eating habits secondary challenges.

## Introduction

In recent years, climate change has intensified as a result of human activities on the environment, leading to an increase in the frequency of extreme weather events, such as heat waves and droughts (1). Food systems play a significant role in climate change, accounting for one-third of global greenhouse gas (GHG) emissions (2) and approximately 70% of global deforestation (3). Extreme weather events also impact food systems, increasing food and nutrition insecurity in vulnerable populations (4). Despite the significant impact of food systems, foods have distinct environmental and health impacts. While grains, fruits, and vegetables have a lower environmental impact, meats generate higher GHG emissions due to their industrial production (5). Replacing animal proteins with plant-based protein sources could reduce cumulative GHG emissions by 14% to 48% (6) and also bring benefits to people’s health (7). For this reason, one of the main strategies encouraged to mitigate the effects of climate change is the reduction of meat production and consumption through the promotion of sustainable diets, such as the flexitarian diet (8,9).

The flexitarian diet is characterized by low meat consumption, particularly red meat (10). This diet could reduce premature mortality related to overweight and obesity by up to 34% (11) and decrease social costs associated with greenhouse gas emissions by between 41% and 74% by 2030 (8), promoting savings of up to 97% in direct and indirect health costs (8). These benefits have spurred an increasing number of studies dedicated to understanding the factors influencing consumption patterns and the reduction of meat consumption among so-called flexitarians (10,12,13). Flexitarians are essentially vegetarians but have occasional meat consumption (10). The voluntary choice for this type of diet, motivated by different individual reasons (e.g., health, convenience) or ethical considerations (e.g., environmental and animal ethics), is the main difference that separates flexitarians from those who reduce meat consumption due to financial constraints (14).

Understanding the motivations and barriers to adopting a flexitarian diet is crucial for promoting sustainable food choices. Food choices play a crucial role in identity formation, often reflecting personal ideologies and life philosophies (15,16). Dietary identity, encompassing how individuals perceive and act towards animal-based products (17), can reinforce sustainable food choices. However, adopting a reduced-meat diet presents several challenges, including attachment to meat’s taste, the cost of alternatives (18), and social environmental factors that influence meat consumption decisions (19,20).

Although scientific literature has explored the barriers to reducing meat consumption, much of this research focuses on developed countries in the Global North (21). Little attention has been given to the development of flexitarianism and its barriers in Global South countries, and few studies discuss flexitarian identity (22,23), creating a gap in understanding the cultural barriers faced in diverse contexts (24). Furthermore, the analysis of motivations and barriers often involves a trade-off between using open-ended questions with smaller samples due to challenges in speech analysis (25) and using closed-ended questions with larger samples (26). From a cultural and sociological perspective (27), considering the novelty of flexitarianism science, it is crucial to gain a comprehensive understanding of the factors influencing the adoption of such diets to develop effective interventions for promoting dietary change. Simultaneously, larger samples can help identify overarching trends.

Therefore, this study addresses these limitations by applying Natural Language Processing (NLP) techniques to a large sample of open-ended responses. This approach allows us to strike a balance between the depth of understanding afforded by qualitative methods and the breadth of insights provided by quantitative approaches. Specifically, we aim to fill this gap by addressing the main challenges associated with reducing meat consumption and adopting a flexitarian diet in Brazil. As a country in the Global South where meat consumption is deeply rooted in the culture (28), Brazil serves as a critical case study for understanding flexitarianism in diverse cultural contexts. By exploring the justifications behind individuals defining themselves as flexitarians and how these self-perceptions contribute to the construction of a flexitarian identity, this research offers insights that can inform global strategies for promoting sustainable diets.

## Materials and Methods

### Study Design and Participants

This is a descriptive and explanatory study utilizing data from a population-based study with a national convenience sample consisting of 1,029 flexitarians in Brazil. Our objective is to identify the main justifications and challenges associated with a flexitarian diet through a mixed analysis (qualitative and quantitative).

We adopted the same criteria and procedures outlined by Teixeira et al. (14) and applied the following inclusion criteria for participants: (i) minimum age of 18 years, (ii) residency in Brazil, (iii) intentional practice of excluding meat from at least one meal per week, and (iv) not being a vegetarian or vegan (see Teixeira et al., 2024 for detailed information on the sampling plan).

### Recruitment of Participants and Data Collection

For data collection, we used an online questionnaire administered from March to June 2022 across Brazil, disseminated through Instagram, WhatsApp, X (formerly Twitter), and Facebook, with the support of volunteers from across the country. At this stage, participants self-reported demographic information, including “gender” (options: cis woman, trans woman, cis man, trans man, non-binary, and prefer not to answer), “ethnicity/color” (options: white, black, yellow, multiethnic, indigenous, and prefer not to answer), and educational level (options: elementary school, high school, undergraduate, master’s, and doctoral).

We also collected information related to the primary source of information about flexitarianism, the main challenges encountered in adopting a flexitarian diet, and the justifications participants consider themselves flexitarians. To this end, we used the following questions: “Why do you consider yourself a flexitarian?” (open-ended), “What was your initial source of information about flexitarianism?” (with options: your parents; school or university; friends; media; other), “What is the main motivation for you to reduce meat consumption?” (with options: concern about environmental impact; animal ethics; health concerns; religion, beliefs, or spirituality; aversion, allergies, or intolerance; other); and “What is the biggest difficulty you face in reducing meat consumption?” (open-ended).

### Data analysis

The data were analyzed using the ChatGPT language model, developed by OpenAI and the Python programming language. For categorical variables derived from multiple-choice responses, such as gender, ethnicity/color, source of information about flexitarianism, and motivations for reducing meat consumption, we used absolute and relative frequencies to summarize the results.

For the analysis of textual data from open-ended questions, we employed Natural Language Processing (NLP) techniques such as Term Frequency-Inverse Document Frequency (TF-IDF) to assess the importance of a word in a document relative to a collection of documents. We then applied matrix factorization techniques, such as Singular Value Decomposition (SVD) and Non-Negative Matrix Factorization (NMF), to identify topics (groups of words) that represent the main themes mentioned by participants. These analytical techniques allowed us to identify recurring themes and response patterns in an automated manner. The following analytical procedures were adopted:

- Data Preprocessing:

The textual data were preprocessed for cleaning and simplification of words. This involved removing stopwords (irrelevant words) and applying stemming (reducing words to their root forms by removing suffixes and prefixes), tokenization (segmenting text into smaller units considering context), and lemmatization (extracting the base form of words). We utilized pipelines to perform these textual data simplification techniques.

- Topic Modeling (BERTopic OpenAI):

We employed the BERTopic (Bidirectional Encoder Representations from Transformers) technique for unsupervised clustering of textual passages. BERTopic uses pre-trained language models that capture semantic and syntactic characteristics of texts. We applied embeddings generated through deep learning techniques via the OpenAI API to BERTopic to enhance topic modeling and improve the performance of the deep learning model in identifying and analyzing topics.

To identify the representative terms for the reasons participants consider themselves flexitarians, we employed the TF-IDF method. We identified 12 topics that presented these reasons. However, some topics generated by the TF-IDF lacked clear motivations.

Therefore, a supplementary human analysis was necessary to define the categories of reasons based on these topics. Through this supplementary analysis, we identified eight categories of justifications (“decreased frequency of consumption,” “concern for health or environment”, “excludes certain animals”, “does not consider meat essential”, “dietary flexibility”, “personal preference”, “social influence,” and “price”), along with two groups that do not present an apparent reason (“no specific justification” and “other”). Subsequently, these categories were grouped into five groups, with titles created based on the keywords of each group: ethical flexitarians, flexible vegetarians, meat appreciators, moderate consumers, and meat reducers.

Next, we applied the c-TF-IDF (Class-based Term Frequency-Inverse Document Frequency) method to analyze the representative terms associated with adherence to flexitarianism, considering the variables of gender, ethnicity, education, and motivations (we differentiate “justification/reason” from “motivation”, understanding that the former relates to participants’ self-perception of what it means to be flexitarian, while the latter refers to the factors that drove this choice). This analysis formed groups of similar documents for each variable. However, the analysis of the reasons mentioned based on demographic and motivational variables revealed an unequal data distribution with low response frequencies (<5), which hindered the application of appropriate statistical tests to assess statistically significant differences between the groups.

For the challenges related to reducing meat consumption and adhering to flexitarianism, we applied the same methods. We used TF-IDF to identify relevant words without stratification by groups, through which we identified three topics of challenges: eating habits, liking the taste of meat, and sources of alternative proteins. We then applied c-TF-IDF to analyze the challenges based on the variables of gender, ethnicity/color, education, and motivation. Unlike the analysis of justifications, the structure of the gender-related data among the challenges was homogeneous, allowing for the analysis of gender differences through the Chi-Square test. However, it was not possible to conduct statistical tests for the variables of motivations, education, and ethnicity/color in this analysis due to the unequal structure of the data.

- Cluster Analysis

We applied cluster analysis to the justifications and challenges related to flexitarianism without stratification and with stratification by groups (gender, ethnicity/color, education, and motivations). We used the K-Means algorithm to select the appropriate number of clusters and visualized the results through a dendrogram generated by complementary hierarchical analysis. To determine the adequate number of clusters, we calculated the silhouette score, which assesses the cohesion and separation of the clusters, indicating the quality of cluster definition. We selected the number of clusters that presented the highest average score.

### Ethical procedures

This research complied with all the regulations governing studies involving human subjects in Brazil as stated in Resolution of the National Health Council No. 466/2012. All research protocols were submitted to the Ethics Committee of the Onofre Lopes University Hospital (CEP/HUOL) at the Federal University of Rio Grande do Norte (UFRN) and were approved under protocol number CAAE 5,348,343. All participants were adults (18 years or older) and provided written informed consent in a designated section of the research questionnaire, with the consent being digitally recorded as part of the questionnaire submission.

## Results

In this study, we analyzed data from 778 participants who met the previously established inclusion criteria. Initially, 1,029 people from all regions of the country participated in the data collection. However, 251 participants were excluded due to the inadequacy of their responses, which were either excessively brief or semantically incoherent concerning the questions about the reasons for identifying as flexitarian and the challenges faced in reducing meat consumption (e.g., “no,” “yes,” “.” etc.), which rendered a detailed textual analysis of their responses unfeasible.

For the presentation of the results, we organized the section as follows: (1) we describe the participants’ profile, including data on gender, ethnicity, education level, and the primary source of information about the flexitarian diet; (2) we highlight the main justifications why participants identify as flexitarians; and (3) we present the main challenges faced in reducing meat consumption and adhering to a flexitarian diet.

### Participant profile and source of information on flexitarianism

Table 1 provides a comprehensive overview of the participants’ demographic characteristics, including gender, ethnicity, educational attainment, as well as their sources of information on flexitarianism and motivations.

**Table 1.**
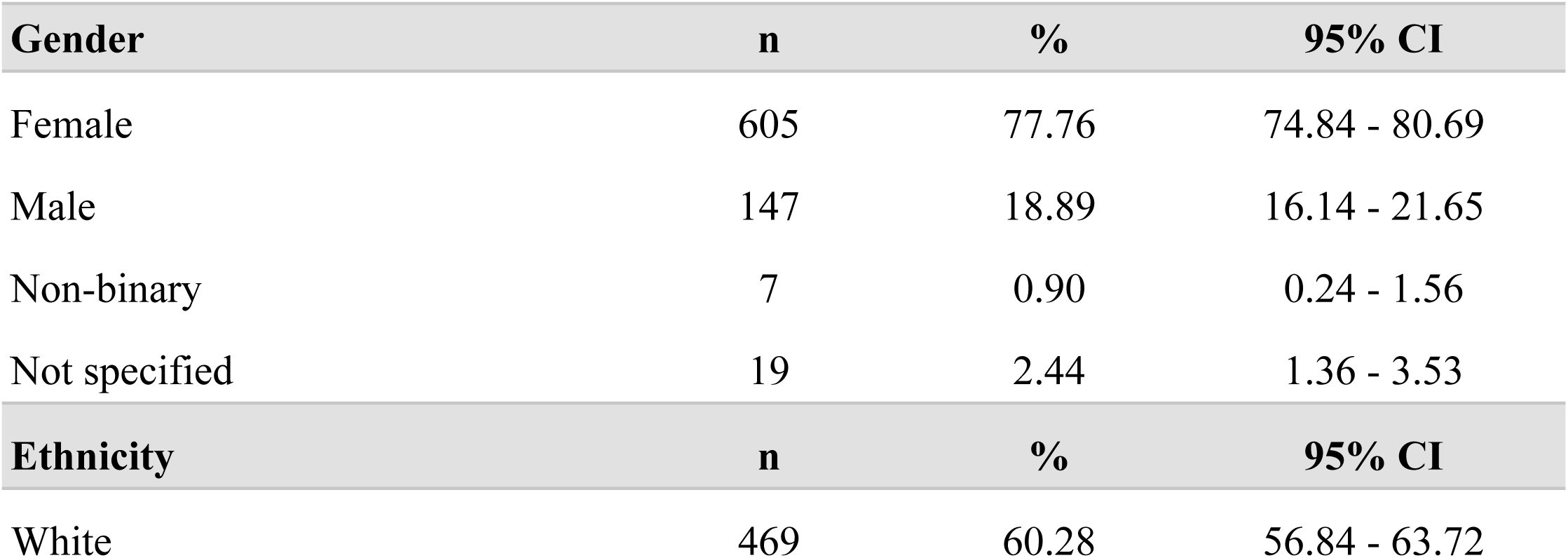
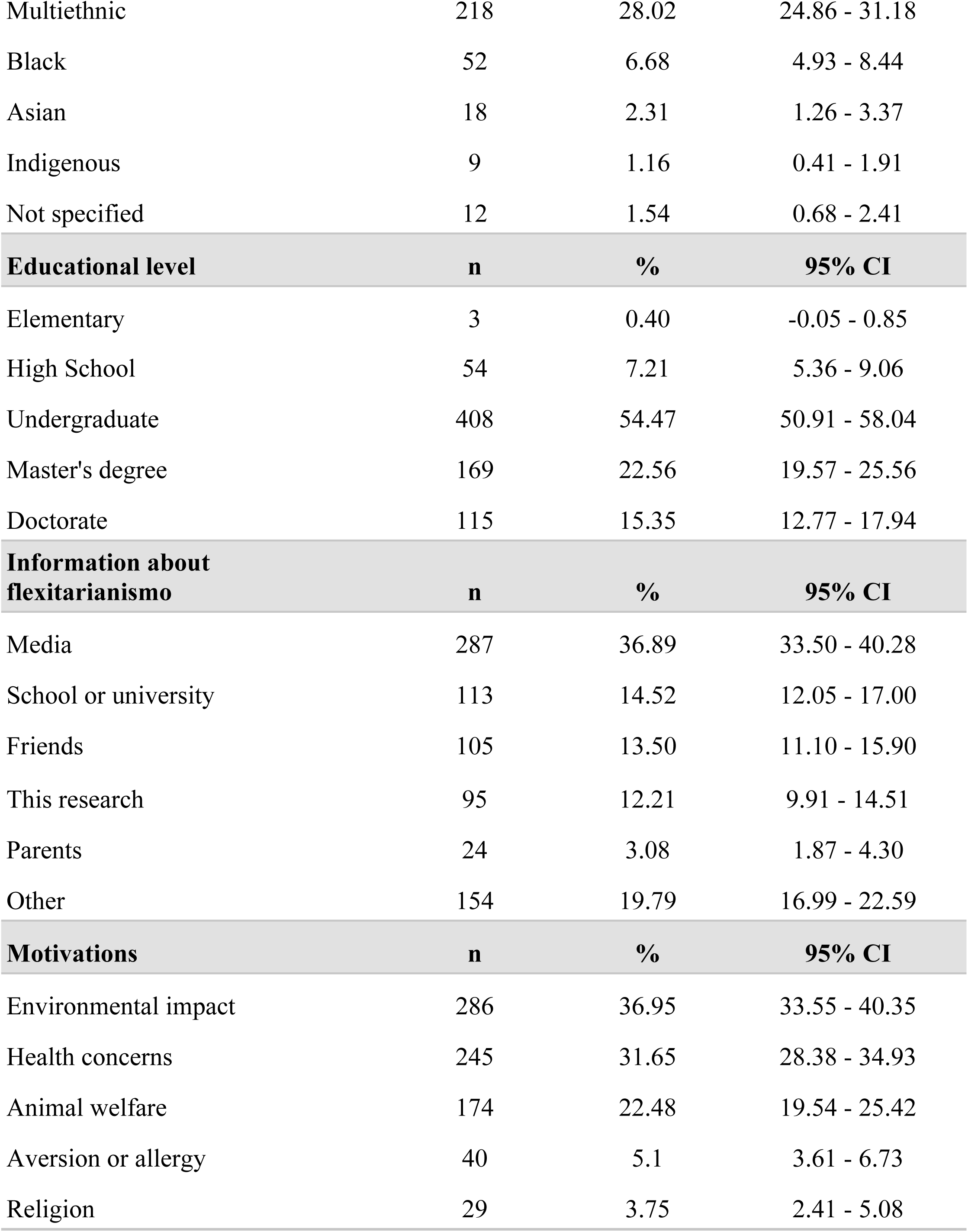
Participant profile, sources of information on flexitarianism, and motivations.

The majority of participants were women with a high level of formal education, indicating a potential link between education, gender, and the adoption of this dietary approach. While the motivations for adopting a flexitarian diet are diverse, two primary themes emerge: environmental concerns and health consciousness. It is also worth noting that while 12% of participants were unfamiliar with the term “flexitarian” before the study, their exposure to this dietary approach stemmed primarily from media sources and educational institutions.

### Why participants considered themselves flexitarians

Our results show that participants identify as flexitarians for a wide range of justifications, grouped into 12 main topics identified through TF-IDF topic modeling. Since some of the topics did not provide clearly defined justifications, we conducted a complementary human analysis and identified eight categories, along with two groups (“no specific justification” and “others”) that do not present an apparent justification. The full list is available in Table 2.

**Table 2.** Justifications for considering oneself as flexitarian.

| Justifications | n | % | 95% CI |
| --- | --- | --- | --- |
| Reduces frequency of consumption | 373 | 47.94 | 44.43 - 51.45 |
| Concern for health or the environment | 253 | 32.52 | 29.23 - 35.81 |
| Excludes some animals | 79 | 10.15 | 8.03 - 12.28 |
| Does not consider meat essential | 19 | 2.44 | 1.36 - 3.53 |
| Flexibility of the diet | 13 | 1.67 | 0.77 - 2.57 |
| Personal preference | 11 | 1.41 | 0.58 - 2.24 |
| No specific justification | 13 | 1.67 | 0.77 - 2.57 |
| Social influence | 6 | 0.77 | 0.16 - 1.39 |
| Price | 5 | 0.64 | 0.08 - 1.20 |
| Other | 6 | 0.77 | 0.16 - 1.39 |

These categories contain examples of full textual excerpts related to each justification for why the participants of the study consider themselves flexitarians. The analysis of the excerpts shows that justifications and motivations sometimes overlap, as in the case of flexitarians, where the justification for identifying as a flexitarian intertwines with motivations related to concerns about health and the environment. To illustrate this, we highlight textual passages from each group of justifications (Box 1).

##### Box 1. Textual passages from group of justifications

**Reduces frequency of consumption:**

*“In recent years, I have reduced the amount of meat I eat at lunch and almost eliminated it at dinner, with weekends being somewhat unpredictable. However, I have no intention of becoming vegetarian or vegan due to the significant planning and sacrifice of personal and social pleasures that this change entails, but I admire the dedication of those who decide to become.”*

**Concern for health or the environment:**

*“Because I became aware of the journey that food takes to reach my plate, and from that, I try to make better choices for my health, the animals, and the planet.”*

**Excludes some animals:**

*“Because I haven’t eaten any red meat or pork for 8 years.”*

**Does not consider meat essential:**

*“I don’t believe it is necessary to consume meat.”*

**Flexibility of the diet:**

*“Because I understand that my way of eating and living does not completely fit into what is understood as vegetarian or vegan, and for me, being flexitarian is the best way to express my lifestyle and food consumption.”*

**Personal preference:**

*“Because I choose not to consume animal products whenever possible.”*

**No specific reason:**

*“I didn’t know this term before the research; I discovered that I am flexitarian now.”*

**Social influence:**

*“Although I want to be a vegetarian, there are factors that prevent me from doing so, and the main one is the meat-centered family culture.”*

**Price:**

*“Price is a mitigating factor, but I have always liked vegetables and legumes in general, and I initially tried to see if I could become vegetarian during the pandemic, and the response was positive. I have maintained my flexitarian lifestyle until today.”*

**Other:**

*“Because I still haven’t managed to completely stop consuming animals.”*

These justifications are part of five distinct groups and are categorized as ethical flexitarians, flexible vegetarians, meat lovers, moderate consumers, and meat reducers. Each group has specific keywords and representative documents identified through topic modeling, using a combination of TF-IDF and BERTopic. One identified topic contains 211 entries deemed irrelevant, grouping terms such as “vegetarian,” “vegetables,” “diet,” and “meat.” The first topic (Topic 1) includes terms such as “diet,” “concern,” “consumption,” and “environmental,” reflecting the participants’ conscious connection between diet and environmental issues. The distribution of keywords (representation) and representative documents for each topic is presented in Table 3.

**Table 3.** Distribution of topic modeling by keywords and representative documents.

| Topic | Name | Representation | Representative Docs | n | % |
| --- | --- | --- | --- | --- | --- |
| 1 | Ethical flexitarians | Food, concern, consumption, environmental, animals | Because I am aware of how animals are raised | 120 | 21.16 |
| 2 | Flexible vegetarians | Vegetarianism, vegetarian, eating, flexitarian | Since I deduce that I no longer consume meat... | 152 | 26.81 |
| 3 | Meat lovers | Meats, taste, want, foods, little | I eat little meat because I don't make a point of it... | 59 | 10.41 |
| 4 | Moderate consumers | Meats, consumption, week, foods | Because there are days of the week when I don't eat meat... | 60 | 10.58 |
| 5 | Meat Reducers | Reducing, reduction, to reduce, reduced, meat | Because I have managed to reduce my meat consumption... | 176 | 31.04 |

These topics follow the same pattern based on gender, with men predominantly in Topic 1 (ethical flexitarians), whose most frequent words are “flexitarian,” “vegetarianism,” “eating,” and “vegetarian.” Women, on the other hand, stand out in Topic 4 (moderate consumers), primarily mentioning words related to reducing meat consumption, such as “reducing,” “reduction,” “reduce,” and “vegetarian.” Individuals with higher education levels (doctorate) indicate environmental issues (Topic 1) as their main reason, using words like “concern,” “meat,” “environmental,” and “food.” Those motivated to adopt flexitarianism due to concerns about environmental impact and animal welfare prevail in Topic 1, while those concerned about health predominately appear in Topic 3 (meat lovers). In Table 4, we present the distribution of topics based on gender, education level, and motivation for adopting flexitarianism.

**Table 4.** Distribution of topics based on gender, education level, and motivation for adoption.

| Gender | Topic 1 |  | Topic 2 |  | Topic 3 |  | Topic 4 |  | Topic 5 |  |
| --- | --- | --- | --- | --- | --- | --- | --- | --- | --- | --- |
|  | n | % | n | % | n | % | n | % | n | % |
| Female | 94 | 78.33 | 115 | 75.66 | 44 | 74.58 | 51 | 85.00 | 139 | 78.98 |
| Male | 22 | 18.33 | 34 | 22.37 | 11 | 18.64 | 7 | 11.67 | 32 | 18.18 |
| Non-binary | 0 | 0 | 2 | 1.32 | 0 | 0 | 0 | 0 | 2 | 1.14 |
| Not specified | 4 | 3.33 | 1 | 0.66 | 4 | 6.78 | 2 | 3.33 | 3 | 1.70 |
|  | **Topic 1** |  | **Topic 2** |  | **Topic 3** |  | **Topic 4** |  | **Topic 5** |  |
| **Educational level** | **n** | **%** | **n** | **%** | **n** | **%** | **n** | **%** | **n** | **%** |
| Elementary | 0 | 0 | 0 | 0 | 1 | 1.69 | 1 | 1.67 | 1 | 0.57 |
| High School | 11 | 9.17 | 17 | 11.18 | 8 | 13.59 | 9 | 15.00 | 14 | 7.95 |
| Undergraduate | 52 | 43.33 | 79 | 51.97 | 34 | 57.63 | 28 | 46.47 | 96 | 54.55 |
| Master's degree | 28 | 23.33 | 35 | 23.03 | 11 | 18.64 | 12 | 20.00 | 39 | 22.16 |
| Doctorate | 29 | 24.17 | 21 | 13.82 | 5 | 8.47 | 10 | 16.67 | 26 | 14.77 |
|  | **Topic 1** |  | **Topic 2** |  | **Topic 3** |  | **Topic 4** |  | **Topic 5** |  |
| **Motivations** | **n** | **%** | **n** | **%** | **n** | **%** | **n** | **%** | **n** | **%** |
| Environmental impact | 53 | 44.17 | 50 | 35.21 | 15 | 25.42 | 26 | 43.33 | 70 | 39.77 |
| Health concerns | 22 | 18.33 | 40 | 28.17 | 23 | 38.98 | 20 | 33.33 | 61 | 34.66 |
| Animal welfare | 38 | 31.67 | 35 | 24.65 | 7 | 11.68 | 10 | 16.67 | 31 | 17.61 |
| Aversion or allergy | 3 | 2.50 | 7 | 4.93 | 13 | 22.03 | 1 | 1.67 | 6 | 3.41 |
| Religion | 4 | 3.33 | 10 | 7.04 | 1 | 1.69 | 3 | 5.00 | 8 | 4.55 |

### Challenges related to adopting flexitarianism

Participants face three main challenges in adopting flexitarianism, according to cluster analysis: eating habits (n = 27, 3.47%), preference for the taste of meat (n = 50, 6.42%), and availability of alternative proteins (n = 702, 90.12%). Analysis of the distribution of relevant words shows that, for the challenge associated with consuming alternative proteins, the most frequently mentioned words, in addition to “eating meat” and “meat consumption,” include “protein source,” “eating at home,” “home option,” as well as additional terms such as “restaurant option,” “substitute meat,” and “find option.” The relevant words associated with each challenge cluster can be seen in Fig 1.

**Fig 1.**
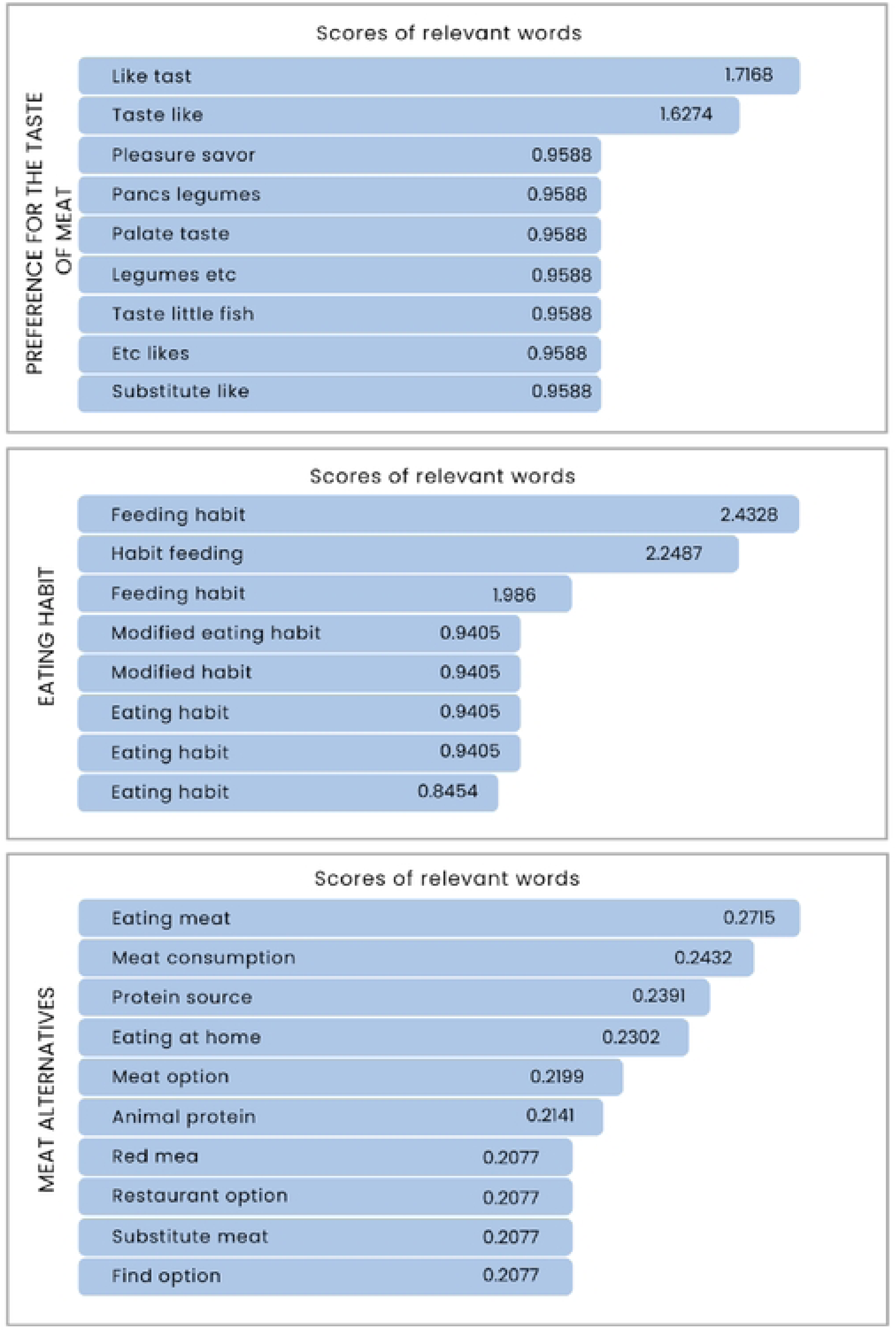
Distribution of relevant words associated with challenges in reducing meat consumption among flexitarians. The figure presents the keywords related to each identified challenge for adopting flexitarianism, along with the corresponding relevance scores for each term as presented through TF-IDF analysis.

Challenges are experienced relatively uniformly among flexitarians across different groups (Table 5). Women face greater barriers related to the taste of meat and dietary habits, while men’s primary challenge is finding alternative options to meat. However, these differences are not statistically significant (χ² = 2.03, p = 0.36).

**Table 5.** Challenges to reducing meat consumption: cluster analysis by gender, education level, ethnicity, and motivation.

| Characteristic | Challenges |  |  |  |  |  |
| --- | --- | --- | --- | --- | --- | --- |
|  | Preference for the taste of meat |  | Eating habit |  | Meat alternatives |  |
| Gender | N | % | n | % | n | % |
| Male | 7 | 4.32 | 4 | 2.47 | 151 | 93.21 |
| Female | 42 | 7.09 | 20 | 3.38 | 530 | 89.53 |
| Educational Level | N | % | n | % | n | % |
| Elementary | 0 | 0 | 0 | 0 | 3 | 100 |
| High School | 3 | 3.66 | 2 | 2.44 | 77 | 93.90 |
| Undergraduate | 24 | 5.83 | 15 | 3.64 | 373 | 90.53 |
| Master's degree | 16 | 9.25 | 2 | 1.16 | 155 | 89.60 |
| Doctorate | 6 | 5.41 | 8 | 7.21 | 97 | 87.39 |
| Ethnicity | N | % | n | % | n | % |
| White | 29 | 6.10 | 14 | 2.94 | 432 | 90.94 |
| Black | 3 | 6.38 | 2 | 4.25 | 42 | 89.36 |
| Indigenous | 0 | 0 | 0 | 0 | 1 | 100 |
| Multiethnic | 14 | 6.54 | 10 | 4.67 | 190 | 88.78 |
| Asian | 3 | 15.00 | 0 | 0 | 17 | 85.00 |
| Not specified | 1 | 7.14 | 1 | 7.14 | 12 | 85.71 |
| Motivations | N | % | n | % | n | % |
| Environmental Impact | 19 | 6.64 | 9 | 3.15 | 258 | 90.21 |
| Health | 14 | 5.86 | 11 | 4.6 | 214 | 89.54 |
| Religion | 2 | 6.25 | 2 | 6.25 | 28 | 87.50 |
| Aversion | 2 | 4.65 | 1 | 2.33 | 40 | 93.02 |
| Animal Ethics | 12 | 6.74 | 5 | 2.81 | 161 | 90.45 |

This trend of minimal variation is similarly observed across participants’ educational levels, with all categories showing a higher percentage in the challenge related to access to alternative options (Table 1). Liking the taste of meat is the most frequent challenge among individuals motivated by animal ethics (6.7%) and environmental concerns (6.64%). Dietary habits are a more common challenge primarily among those motivated by religion or spirituality (6.25%). Access to alternative protein options affects all motivations uniformly, but it is particularly significant for those who have a strong aversion to meat (93%).

## Discussion

This study aimed to identify the primary justifications and challenges faced by Brazilian flexitarians, providing significant contributions to understanding the dynamics and challenges associated with a flexitarian diet. In summary, we highlight two main findings: (i) we identified the absence of a homogeneous flexitarian identity, distributed across five distinct clusters: ethical flexitarians, flexible vegetarians, meat lovers, moderate consumers, and meat reducers; and (ii) we found that the primary challenge faced by our participants is directly related to the social environment, specifically the difficulty of finding alternatives to meat, which makes cultural issues secondary.

### Flexitarian identity: why participants consider themselves flexitarians

We identified eight categories of justifications for why participants consider themselves flexitarians. These justifications are directly related to both facilitating factors for adopting flexitarianism (not considering meat essential for nutrition, dietary flexibility, and personal preferences for reducing meat consumption) and the challenges faced in this adoption (social influence/unfavorable environment for reducing meat consumption and the cost of meat and a sustainable diet). The eight categories fit into five groups of flexitarians: ethical flexitarians, flexible vegetarians, meat lovers, moderate consumers, and meat reducers.

We refer to ethical flexitarians as those who express a greater concern for the environment and animals, seeking to minimize the impact of their food choices. Flexible vegetarians are those who often express a desire to become vegetarian but still consume meat for various reasons, ranging from a meat-based food culture to a lack of the necessary discipline to complete the transition. On the other hand, meat lovers, moderate consumers, and meat reducers are more closely related, as they tend to reduce meat consumption due to the perception that meat is not necessary in their diet, although they still enjoy its taste. These findings align with Randers & Thøgersen’s (22) definition of flexitarian identity, which emphasizes an awareness of the negative consequences of excessive meat consumption while balancing these concerns with other goals and priorities. This identity is also influenced by the social environment, with those in socially supportive settings for meat reduction endorsing this behavior more strongly (22).

The concept of dietary identity, which refers to individuals’ self-perception regarding their consumption and avoidance of animal-derived products (29), enables the interpretation of discrepancies between dietary patterns and diet labels. Thus, adhering to a flexitarian dietary pattern (i.e., reducing meat consumption) and self-identifying as flexitarian may diverge. For example, we found that many participants were unfamiliar with the flexitarian diet, reporting that their first encounter with the term occurred during this research. This finding may suggest (i) limited familiarity with the terms “flexitarian” and “flexitarianism”, possibly due to the still-nascent promotion of this diet even among its followers, and (ii) that the analyzed flexitarians do not fully express the social identity surrounding their diet, as they occupy a continuum between identifying as omnivores or vegetarians (23). In analyzing the centrality of a meat-reduced diet to their identity and beliefs about carnism (the ideology of eating animals), (30) found that flexitarians identify more strongly as vegetarians. This flexitarian identity remains an emerging and weakly consolidated trait among our participants, suggesting that many do not recognize themselves as part of the flexitarian group. This lack of identification may hinder both adherence to and maintenance of this diet due to the absence of a sense of belonging to a social group. Greater promotion of the benefits of a flexitarian diet, as well as strengthening the identity associated with it, may serve as effective strategies for encouraging the adoption of more sustainable diets.

### Challenges in reducing meat consumption and adopting flexitarianism

Among the main challenges to following a flexitarian diet among our participants are aspects related to the taste of meat, eating habits associated with its consumption, and the unavailability of alternative options. This finding aligns with barriers identified in various studies (18,31,32), which highlight sensory appeal, specifically the attachment to the taste of meat, and the scarcity of plant protein options in markets and restaurants as primary challenges to reducing meat consumption. While finding alternatives to meat is the main challenge faced by most of our participants, we believe this barrier can be minimized through interventions that encourage and enhance nutritional knowledge about adequate food alternatives and their preparation (25), as a lack of culinary skills is a significant barrier to adopting plant-based diets (33). Furthermore, the development of plant proteins that mimic the taste of meat can help overcome this challenge, given that liking the taste of meat is an important factor for consumers (34). However, this approach should be interpreted cautiously, as we lack clarity on the effects of producing and consuming plant-based foods on human health and the environment.

We believe that the challenges related to the enjoyment of meat and eating habits share a common characteristic that complicates overcoming these barriers: the familiarity and availability of food shaped by culture. Repeated exposure to a food item, such as meat, tends to increase preference for it (35). Thus, familiarity influences people’s food preferences and the importance they attach to its taste and consumption (35,36). Cultural traditions and family practices not only influence exposure patterns to foods but also play a significant role in selecting appropriate foods for special occasions and celebrations (35,36). For instance, meat consumption tends to be higher during group meals with family or friends and in outdoor settings, such as restaurants (20). This is because the social environment is one of the main factors driving increased meat consumption (18). In this context, social norms, connections, and social networks can significantly influence the decision to reduce or consume meat (19,20). Support from social networks (friends, family, partners, etc.) that encourage the reduction or elimination of meat consumption is also an aspect that can influence individuals’ decisions to decrease their meat intake (31). For this reason, we believe that effective interventions to overcome these challenges should focus not only on increasing the availability of plant-based alternatives to meat but also on reconfiguring the sociocultural role that meat plays, which can be achieved through media and educational strategies.

Moreover, other factors, such as the cost of a sustainable diet, pose important challenges to consider, as the price of food was mentioned as a factor by five participants. This is a significant barrier that must be taken into account both when introducing plant-ble diet, such as the flexitarian diet, can cost up to 60% more than a convased products to the market and in guidelines for reducing meat consumption, as a sustainabentional diet (8). Although reducing and completely eliminating meat can yield significant benefits for the planet, meat plays a complex role in people’s nutrition and culture (28,37). Its exclusion may have adverse effects on individuals’ health and deepen existing inequalities among those whose income and diets depend on this source, particularly in countries in the Global South (38). Therefore, it is essential to consider the role of meat consumption in different regions and cultures, as well as to provide means to support the dietary transition towards greater consumption of biodiverse foods, without compromising people’s right to adequate nutrition (8).

### Limitations

Among the main limitations of this study are: (i) the use of a non-probabilistic sample, which restricts the generalization of our conclusions; (ii) the lack of awareness about the flexitarian diet among a considerable number of our participants; (iii) the absence of data on the culinary and nutritional knowledge of the participants, as these factors could pose challenges for a diet with reduced meat intake; and (iv) the lack of inferential analyses: the unequal structure of the data among demographic and motivational groups prevented the execution of inferential tests, such as the Chi-square test and Fisher’s Exact Test, to determine the statistical significance of the differences between the groups. This occurred because, in addition to the observed frequency being less than the expected (<5), the categories of reasons and challenges within each group exhibited inequality in the number of observations. We considered aggregating categories; however, we opted to report the observed differences without conducting inferential analyses to maintain the descriptive nature of our study. Furthermore, we did not extrapolate our results beyond the analyzed sample and believe that allowing participants to self-report their own challenges and reasons, without imposing pre-established categories, may have contributed to mitigating these limitations, particularly the explanatory ones regarding the actual reasons for considering themselves flexitarians. As this research focused on the challenges and self-concept related to identifying as a flexitarian, we did not assess the predictive potential of psychological traits.

## Conclusion

This study offers significant contributions to the understanding of flexitarian diets, using Brazil as a case study to explore broader global trends in developing countries. By employing a mixed-methods approach enhanced by Natural Language Processing, we identified five distinct clusters of flexitarian identity. These categories illustrate the diverse motivations and self-perceptions that characterize flexitarianism, reflecting a complex interplay of ethical, health, and personal factors. Our findings highlight the pivotal role of the social environment in influencing dietary choices, with the primary challenge being the limited availability of meat alternatives. This barrier, which surpasses cultural influences, suggests a need for targeted interventions to enhance access to affordable, sustainable food options. Addressing this challenge is crucial for facilitating the transition to flexitarian diets, particularly in regions where meat consumption is deeply embedded in cultural practices. Furthermore, the study reveals that demographic variables, such as gender, do not significantly affect the challenges faced by flexitarians, indicating a potential uniformity in the barriers encountered across different groups. This insight is valuable for policymakers and the food industry as they craft strategies to promote sustainable dietary practices. By examining Brazil, a country with a rich cultural tapestry and significant meat consumption, this research provides a lens through which to understand flexitarianism in diverse contexts. Future research should expand on these findings by exploring regional and cultural variations in flexitarian adoption, particularly in other Global South countries. Such studies will deepen our understanding of the global dynamics of this dietary trend and inform strategies to support the broader adoption of flexitarian diets, contributing to environmental sustainability and public health worldwide.

## Acknowledgments

We would like to thank all volunteer researchers who helped publicize the data collection instrument.

## Author contributions

Conceptualization: CDT, SMG, MCMJ; Data curation: CDT; Formal analysis: EJMN, CDT; Investigation: CDT; Methodology: CDT, SMG, MCMJ; Supervision: MCMJ; Writing - original draft: CDT; and Writing - review & editing: DGL, SMG, MCMJ.

